# Transcriptional Profiles Analysis of COVID-19 and Malaria Patients Reveals Potential Biomarkers in Children

**DOI:** 10.1101/2022.06.30.498338

**Authors:** Nzungize Lambert, Jonas A. Kengne-Ouafo, Wesonga Makokha Rissy, Umuhoza Diane, Ken Murithi, Peter Kimani, Olaitan I. Awe, Allissa Dillman

**Affiliations:** Liverpool School of Tropical Medicine Research Unit, Centre for Research in Infectious Diseases (CRID), P.O. Box 13591, Cameroon; African Institute of biomedical science and technology (AiBST), Wilkins Hospital, Block C, Corner J. Tongogara and R. Zimbabwe; University of Rwanda, Department of Biology, Rwanda; Synthetic Biology and Omics Data Center, SynbioRwanda, Rwanda; Chinhoyi University of Technology (CUT), P.BAG 7724, Zimbabwe; International Centre of Insect Physiology and Ecology (ICIPE) P.O. Box 30772-00100, Kenya; University of Ibadan, Ibadan, Nigeria; National Institutes of Health, Bethesda, MD, U.S.A.

**Author notes:** Corresponding author: Nzungize Lambert.

**Keywords:** Malaria, COVID-19, children, RNA seq, lncRNA, gene expression, biomarker

## Abstract

The clinical presentation overlap between malaria and COVID-19 poses special challenges for rapid diagnosis in febrile children. In this study, we collected RNA-seq data of children with malaria and COVID-19 infection from the public databases as raw data in fastq format paired end files. A group of six, five and two biological replicates of malaria, COVID-19 and healthy donors respectively were used for the study. We conducted differential gene expression analysis to visualize differences in the expression profiles. Using edgeR, we explored particularly gene expression levels in different phenotype groups and found that 1084 genes and 2495 genes were differentially expressed in the malaria samples and COVID-19 samples respectively when compared to healthy controls. The highly expressed gene in the COVID-19 group we found CD151 gene which is facilitates in T cell proliferation, while in the malaria group, among the highly expressed gene we identified GBP5 gene which involved in inflammatory response and response to bacterium. By comparing both malaria and COVID-19 infections, the overlap of 62 differentially expressed genes patterns were identified. Among them, three genes (ENSG00000234998, H2AC19 and TXNDC5) were highly upregulated in both infections. Strikingly, we observed 13 genes such as HBQ1, HBM, SLC7A5, SERINC2, ATP6V0C, ST6GALNAC4, RAD23A, PNPLA2, GAS2L1, TMEM86B, SLC6A8, UBALD1, RNF187 were downregulated in children with malaria and uniquely upregulated in children with COVID-19, thus may be further validated as potential biomarkers to delineate COVID-19 from malaria-related febrile infection. The hemoglobin complexes and lipid metabolism biological pathways are highly expressed in both infections. Our study provided new insights for further investigation of the biological pattern in hosts with malaria and COVID-19 coinfection.

## Introduction

The clinical diagnosis and distinction between malaria infection and COVID-19 in children presenting with malaria symptoms at a health care facility is a challenge to clinicians due to their overlapping symptoms. This causes a potential risk of misdiagnosis and in turn inappropriate treatment, therapy provision or untimely preventable death. The age structure and demography play a key role in COVID-19 mortality, where death tends more in elders than children [1]. Malaria is an opportunistic parasitic infection documented to be the leading cause of mortality and morbidity globally [2]. Sub-Saharan Africa bears the highest burden of the disease, with *Plasmodium falciparum* contributing to the most severe form of the disease. Over the past two decades, the aversion of 1.5 billion malaria cases and 7.6 million malaria related deaths, has been greatly attributed to the use of Long-lasting Insecticidal nets (LLINs), Indoor residual spraying (IRS), and Artesunate combination therapy (ACT) as rolled out by existing national malaria control programs (NMCPs) [3]. Nonetheless, the recent WHO report still documents 229 million new infections and 409000 deaths globally as of 2019, with the highest observed mortality occurring in children under the age of five [4–6]. In this vulnerable population, malaria is quite severe and leads to the majority of most hospital admissions. In 2020, due to the disruption to service because of the COVID-19 pandemic, the malaria case incidence increased in Africa and counted about 95% of cases.[3]

COVID-19 disease is caused by a novel coronavirus, severe acute respiratory syndrome coronavirus 2 (SARS-CoV-2), in Wuhan, China in 2019, and subsequent spread globally [7]. To date, there is no specific treatment for COVID-19 despite the volume of information published in different renowned journals [8, 9]. Moreover, the nature of the disease coupled with overlapping symptoms brought with it a lot of confusion, especially in Sub-Saharan Africa where the burden of malaria alongside other infectious diseases is high. Recently, a lot of clinical studies have documented a strong relationship between severe malaria infection and SARS-CoV-2 in adults across the globe [10–12]. Pathogen diagnosis depends to a great extent on the interaction between the host and pathogen which usually leads to the production of detectable biomolecules either from the pathogen or the host that can be used to detect the presence of the pathogen.

Over the years, standard laboratory techniques used in the study of the role of genes in disease development include Northern blotting [13, 14], which allow the study of gene expression via RNA detection [15]; quantitative PCR [16, 17], which contributes to the detection, characterization and quantification of RNA transcripts; and Microarray analyses, used to simultaneously detect the expression levels of multiple genes at a time. Nonetheless, their limitation is the requirement of prior knowledge of genes, transcripts and the availability of a limited number of probes. Recently, RNA seq gene expression profiling using Next-generation sequencing (NGS) has supplemented microarrays as the preferred method for transcriptome wide identification of DEGs (differentially expressed genes) [18, 19]. Like other NGS platforms, this technique allows for massive high throughput sequencing, identification of novel transcriptomes, and the ability to perform single nucleotide resolution.

The advances in bioinformatics technology have enabled subsequent downstream analyses of the sequencing platform outputs. This includes the provision of high-quality visual outputs using qualitative and quantitative data that clearly describe what is going on at the transcriptomic level [20–22]. These are inclusive of principal component analysis plots (PCA), sample-to-sample distance plots, dispersion estimate plots, histogram of *P*-values plots, and MA plots. Recently, enrichment analysis tool kits provided visualization of overexpressed genes and transcripts in the treatment and affected groups, including the pathways associated with the observed differences [23–25]. This study provides a tool that utilizes RNA seq data for the identification of a biomarker that can help in the accurate and timely diagnosis of COVID-19 infection in children presenting with severe malaria symptoms and vice versa (Table 1).

**Table 1.** Similarities in clinical and pathophysiological presentations between malaria and COVID-19 infections.

|  | **Malaria** | **COVID-19** |
| --- | --- | --- |
| Incubation period | Varies from 7 to 30 days<br>(Average 7-14 days). | Range from 2 to 14 days<br>(Average 5-6 days). |
| Pathophysiology | A cytokine storm triggers an exaggerated inflammatory response that may damage blood vessels, kidneys, the liver and lungs. |  |
|  | Acute respiratory distress syndrome (ARDS) from pulmonary thrombosis consequent to cytokine storm. |  |
| Fatality | Fatal in children and pregnant women. | Mild in children and Fatal in elderly |
| Symptoms | Chest retraction | Trouble breathing |
|  | Impaired taste | Loss of smell and taste |
|  | Diarrhea, vomiting and Cough (frequent in children with malaria). | Diarrhea, vomiting and Cough. |
|  | Headache, muscle or joint pain. |  |
|  | Fever, High temperature 38 °C or above |  |

The devastating impact of infectious disease outbreaks and pandemics on health systems could be overwhelming, especially when there is an overlap in clinical presentations with other disease conditions, for instance cases of malaria and COVID-19 (Table 2). We hypothesized that there can be a biomarker based on the children’s immune responses against both infections. Therefore, by using RNA seq datasets available on open access databases, we explored the biological signatures and further characterized the distinctiveness of each etiological presentation classification at the transcriptome level.

**Table 2.** Prevalence of children exposure to malaria and COVID-19 infections.

| Condition | Prevalence (%) | Region | Ref. |
| --- | --- | --- | --- |
| Malaria | 16 | Global | [27] |
| COVID-19 | 18.9% | United States | [28] |
| Malaria and COVID-19 coinfection | 11 | Global | [29] |

## Materials and Methods

RNA seq data paired end reads produced by NGS platforms (Illumina HiSeq 2500 and Illumina NovaSeq 6000) from public repositories such as Gene Expression Omnibus (GEO) and European Nucleotide Archive (ENA) were used for the study. The reference genome was obtained from open-source databases such as ENSEMBL and <u>GENCODE</u>. All RNA seq datasets had been designed for transcriptomic analyses.

### Data collection

The RNA-seq datasets were collected to study how the gene expression fluctuates under the host responses to pathogens. Specifically, our study examined the host responses profile using gene expression levels in COVID-19 and malaria patients (Fig. 1). As depicted on figure 1A, keywords such as children, fever, malaria, COVID-19, pediatrics, peripheral blood etc. were used to search and download malaria, COVID-19 related transcriptome data sets from Expression Omnibus (GEO) and European Nucleotide Archive (ENA) databases. The data was downloaded under the fastq format as raw data.

**Figure 1.**
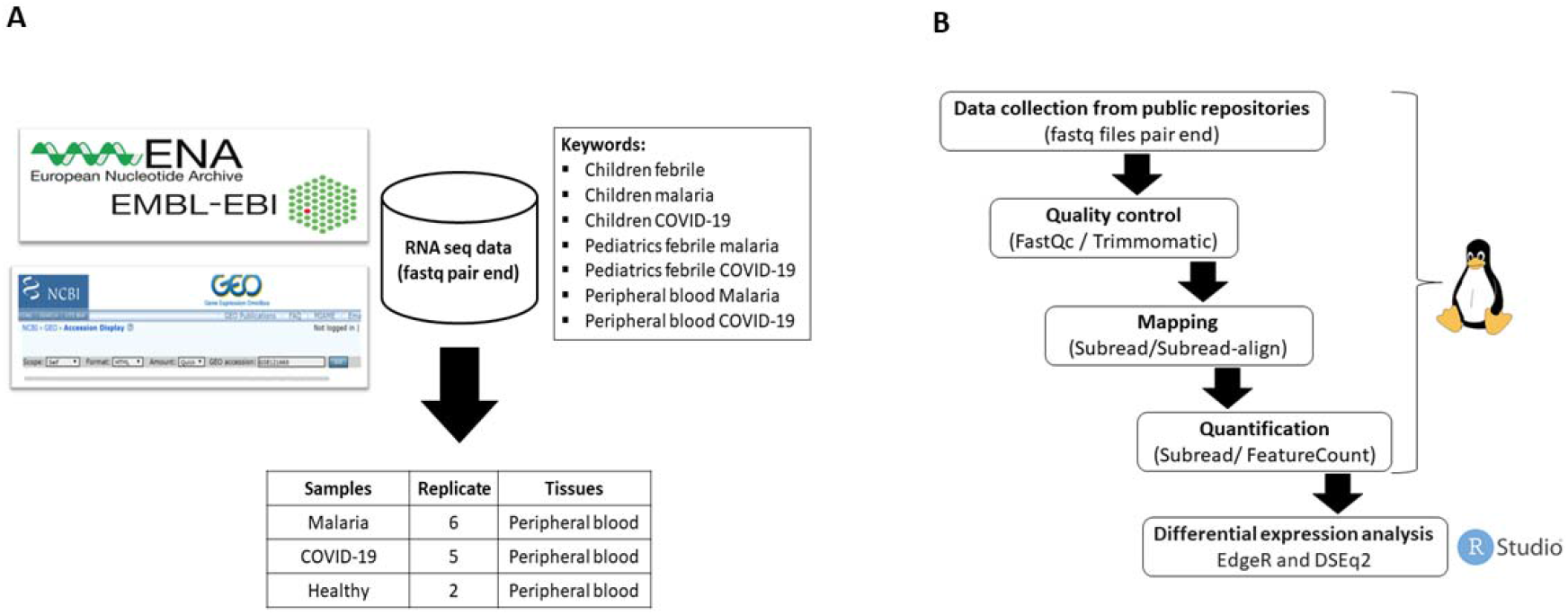
An illustration of the experimental design and RNA seq data analysis. **A)** A total of 13 RNA seq datasets of infected children were collected from public repositories, categorized into three groups: healthy as a control (2 samples); COVID-19 patients (5 samples); and malaria patients (6 samples). **B)** Downstream analysis, a workflow demonstrates the preprocessing of raw sequence data on the Linux platform, as well as downstream analysis of RNA-seq data using the R packages under the RStudio platform.

### Data preprocessing and mapping

The downloaded raw data fastq files made up of paired end reads of between 100-150 bp in length were generated from 13 samples categorized into three groups; a group of COVID-19 samples with five biological replicates, a group of malaria samples with six biological replicates and a healthy group considered as control with two biological replicates. The initial processing and quality assessment of the raw sequence reads was performed using FASTQC (v0.11.9, Babraham Bioinformatics, UK) to identify good reads with default parameters. Adapter trimming was then performed using the Trimmomatic (v0.39) tool [26]. Trimming was done using the following parameters LEADING:3 TRAILING:3 SLIDINGWINDOW: 4:15 MINLEN:36) to remove bad reads and low-quality reads. The cleaned reads were aligned to the human reference genome downloaded from the Gencode database (GRCh38.primary_assembly.genome.fa) and the annotation file was downloaded from the ENSEMBL database (Homo_sapiens.GRCh38.104.gtf). Therefore, a reference genome was indexed with Subread/subread-build index tool followed by the alignment of cleaned reads to the human reference genome using Subread/Sub-read align tool. Afterwards, transcript quantification was performed using the featureCounts option of the Subread package (v.2.0.1). The generated matrix was then used as input for downstream analysis to determine of differentially expressed genes using R packages (Fig.1B).

### Differential gene expression analysis

For differential expression analysis, edgeR (v3.34.0) a Bioconductor package was used in the R software (v4.1.0) to determine the expression difference between groups. To ensure that all samples had a similar range and distribution of expression values, we performed a normalization process using a TMM (Trimmed Mean of M-values) method. This method was preferred because it accounts for sequencing depth, RNA composition, and gene length. It is also appropriate for comparisons between and within samples. We used PCA (principal component analysis) to visualize the distribution of RNA seq data between and within groups. In edgeR, Benjamini and Hochberg’s method was used for estimating the adjusted p-values. In our study, by using a cutoff of 0.05 no significantly expressed genes were detected upon COVID-19 and healthy samples comparison. Therefore, to continue with the analysis, the adjusted p-values were relaxed to 0.1 to get the subjective significant expressed genes. We considered all genes with an adjusted *p* value (padj< 0.1) as significant (a fraction of 10% false positives acceptable) and differentially expressed genes with fold change (absolute log2) of 1 (abs(log2FoldChange) > 1). The DAVID-Functional Annotation Tool was used for functional enrichment analysis based on the differentially expressed genes between both infection groups (COVID-19 and malaria) and the GOplot 1.0.2 package used for visualization. K-means clustering was performed using the Integrated Differential Expression and Pathway analysis 95 (iDEP-95) platform (http://bioinformatics.sdstate.edu/idep/) to determine the differentially expressed genes between the different groups (Malaria and COVID-19).

## Results

### RNA seq datasets characteristics

Preliminary analysis showed a disparity in library sizes between groups (Fig. 2A) hence the normalization process (Fig. 2B). Following PCA using normalized expression data, samples grouped distinctively by clinical status with the COVID-19 samples showing a marked level of scattering (Fig. 2C). We explored the correlation in term of gene expression between our study groups. As shown with the PCA, there was a great correlation between samples of the same clinical group indicating similar gene expression in malaria, COVID-19 and healthy as a control. (Fig. 3).

**Figure 2.**
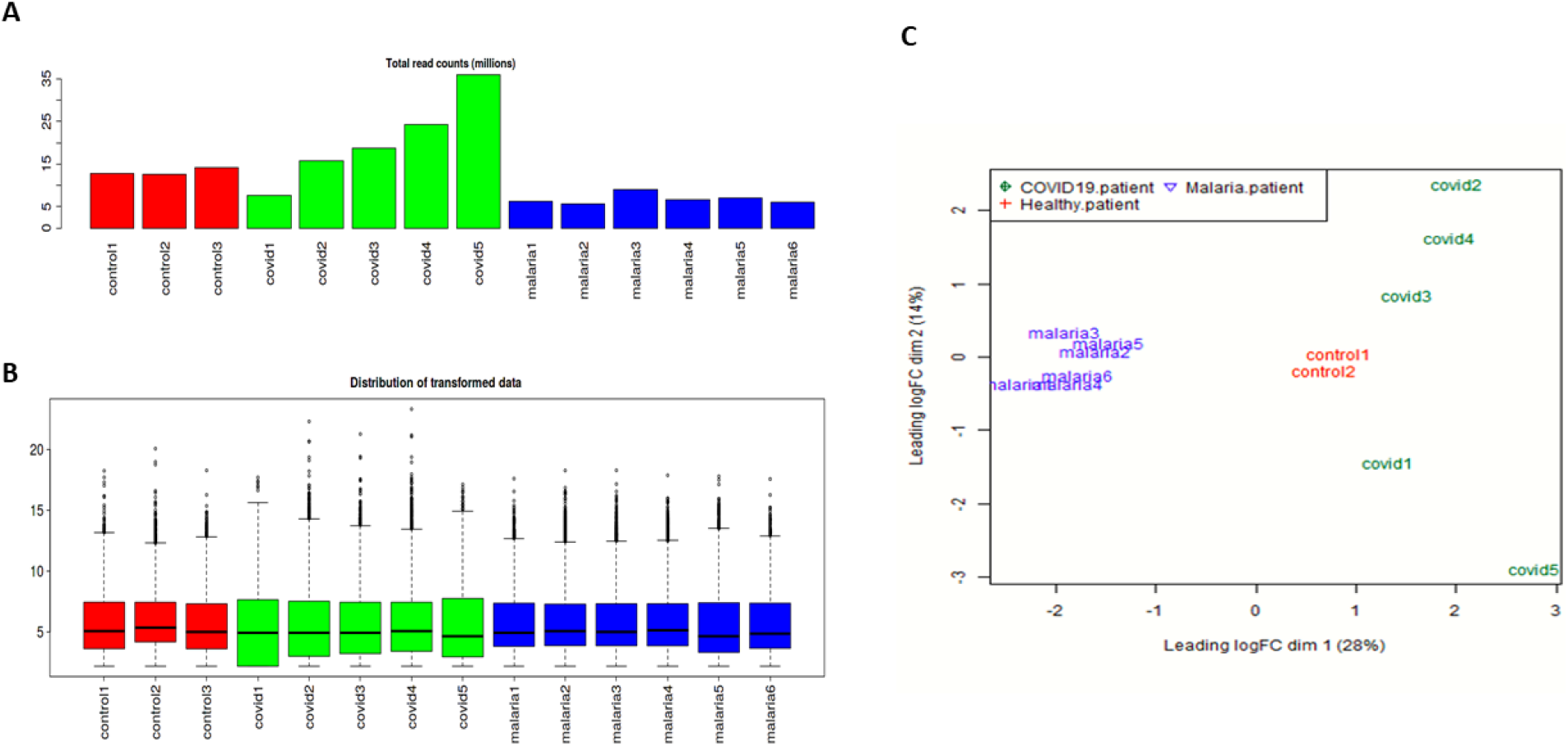
Distribution of RNA seq data between samples. **A)** Before normalization. **B)** After normalization. **C)** Principal component analysis (PCA) of children transcriptome from PBMC of different conditions: children infected with COVID-19, children with malaria infection, and healthy children. COVID-19 samples and malaria samples were clearly separated. However, within the COVID-19 group, samples were relatively scattered, indicating poor consistency.

**Figure 3.**
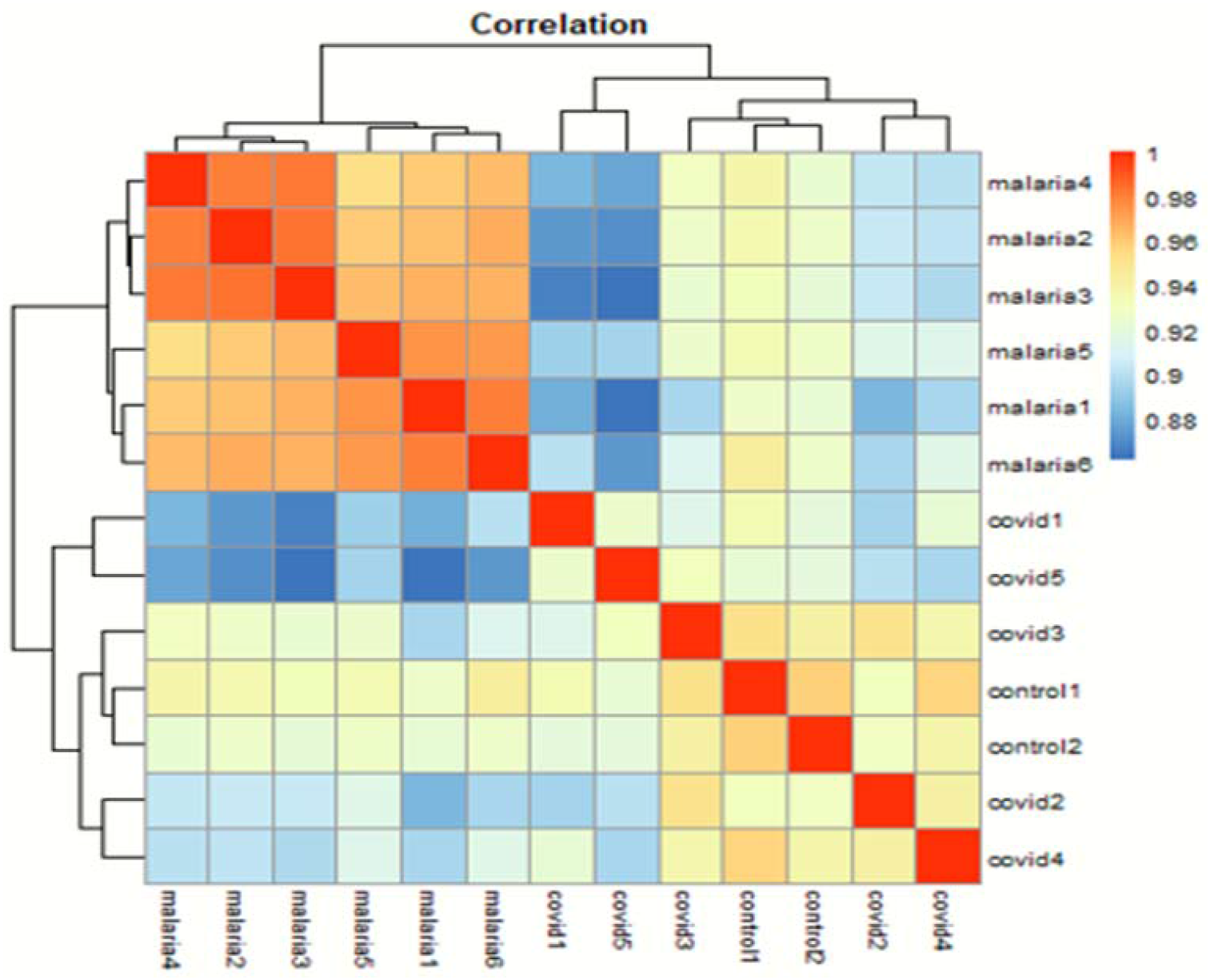
Heatmap comparing the relative expression levels of genes differentially expressed between children with of COVID-19 and malaria infection versus the healthy donor.

### COVID-19 patients expressed genes that are associated with signaling and virus-induced tumours

To assess transcriptional level between children infected with COVID-19 and healthy, a differential expression analysis (DEA) was performed and the outputs visualized with a volcano plot while considering an abs fold change > 1 and adjusted p-value (padj) value < 0.1 as significant. Therefore, DEA revealed 2495 genes differentially expressed between children with COVID-19 and healthy controls (188 upregulated and 2307 downregulated) (Fig. 4A). The top up-regulated genes were found to be likely to play a role in cell signaling and or ion transport across the cell membrane, humoral immune response. These include the TCAF2C, METTL7B, AChE and HBQ1. In addition to signaling, genes associated with asthma, flu (influenza), virus-induced tumors, lung and cervical cancers. Wong and Thai were also found among the top upregulated ones such as Rap1GAP, CD151, UBE2C and NECTIN2 [30]. MAP2K3 gene (mitogen-activated protein kinase 3) based on the previously published data, has been proposed as a potential drug target due to its activity in lung inflammatory processes (Table 3) [31]. We performed GO (gene ontology) annotation analysis to explore the biological functions of the DEGs in COVID-19 group (Fig. 4B). The analysis revealed that the DEGs were significantly associated with signal transduction. The following processes were found enriched in regulation of membrane potential, regulation of post-synaptic membrane potential and sensory perception of smell, ion transport and cell-cell adhesion among others (Fig. 4B).

**Figure 4.**
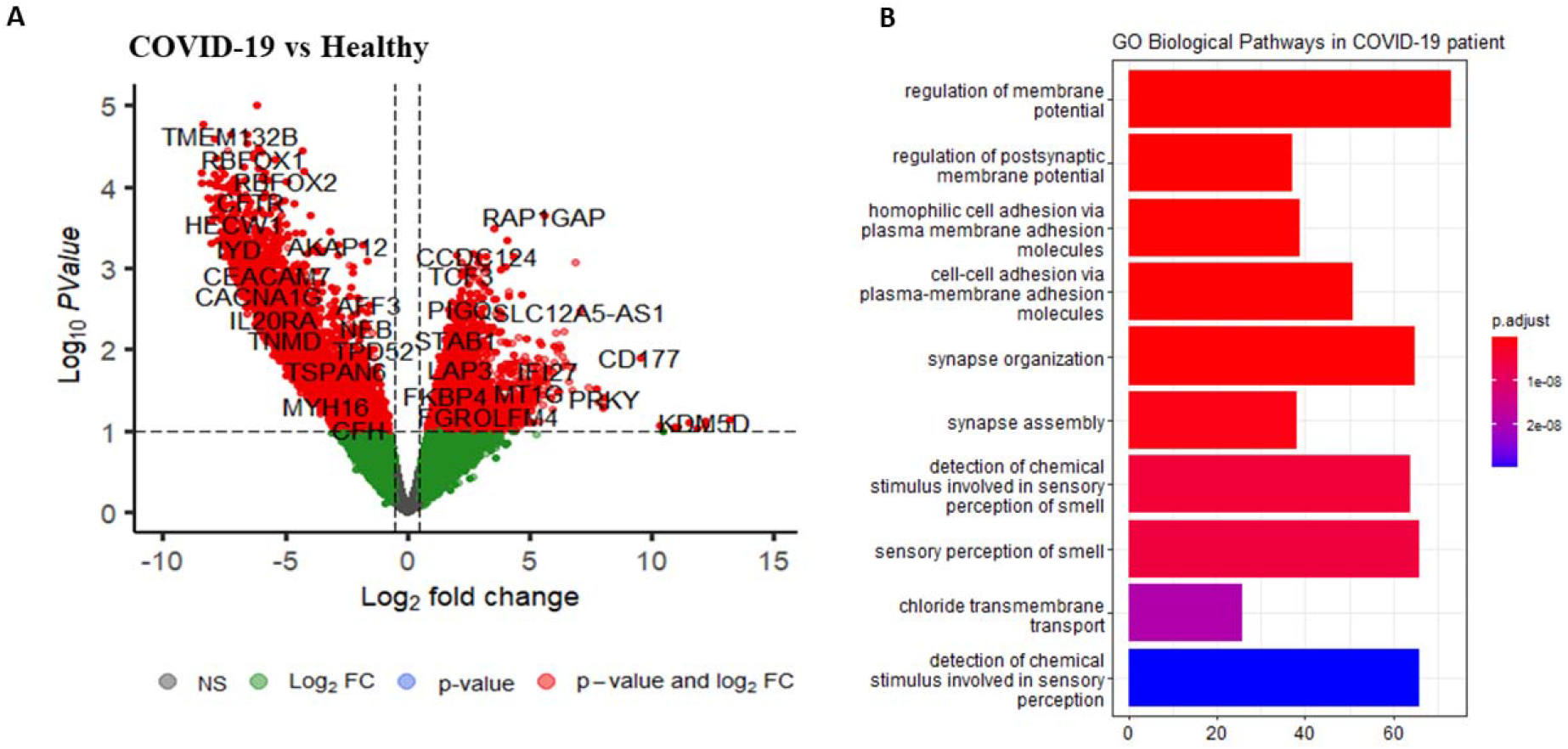
Volcano plot comparing COVID-19 infected individuals and healthy. The X-axis shows log□ fold change (positive values are up regulated relative to healthy). The Y-axis shows the −log10 of BH adjusted p-value (padj) value. The horizontal dashed line marks P= 1%, and the vertical dashed lines indicate log2 fold expression difference among conditions. The differentially expressed genes are indicated in red according to a P<.01 and a log2FC > 1, green points represent genes passing the log2FC threshold but not the adj. p value, and grey points represent genes with no significant difference for both thresholds.

**Table 3.** List of top ten selected genes differentially expressed in children with COVID-19.

| Gene name | Description | logFC | padj |
| --- | --- | --- | --- |
| SLC12A5 | Solute Carrier Family 12 Member 5 | 7.0 | 0.0782 |
| GPS2P1 | G Protein Pathway Suppressor 2 Pseudogene 1 | 6.4 | 0.0909 |
| RCCD1-AS1 | RCC1 Domain Containing 1 | 6.0 | 0.0934 |
| RAP1GAP | RAP1 GTPase Activating Protein | 5.5 | 0.0652 |
| METTL7B | Methyltransferase Like 7B | 5.4 | 0.0997 |
| C1QC | Complement C1q C Chain | 4.9 | 0.0985 |
| MRPL23 | Mitochondrial Ribosomal Protein L23 | 4.6 | 0.0717 |
| TCAF2C | TRPM8 Channel Associated Factor 2C | 4.5 | 0.0963 |
| ITGA7 | Integrin Subunit Alpha 7 | 4.3 | 0.0652 |
| HBQ1 | Hemoglobin Subunit Theta 1 | 4.1 | 0.0714 |
| ACHE | Acetylcholinesterase | 4.1 | 0.0717 |
| CD151 | CD151 Molecule | 4.0 | 0.0652 |
| DCTN6-DT | DCTN6 Divergent Transcript | 4.0 | 0.0999 |
| WASF1 | WASP Family Member 1 | 3.9 | 0.0667 |
| UBE2C | Ubiquitin Conjugating Enzyme E2 C | 3.8 | 0.0667 |
| NECTIN2 | Nectin Cell Adhesion Molecule 2 | 3.8 | 0.0787 |
| MAP2K3 | Mitogen-activated protein kinase kinase 3 | 2.5 | 0.0752 |

logFC: Log2fold change; padj: adjusted *P* value

### Malaria patients expressed genes indicative of vacuolar lumen interactions

Next, we sought to compare children with malaria and the healthy controls, we found 1084 (179 upregulated and 905 down regulated) gene differentially expressed (Fig. 5A). Based on the DEGs analysis, we found the following GO-term associated with vacuolar membrane, lysosomal lumen, lytic vacuole membrane, histone methyltransferase complex to be significantly enriched in malaria group (Fig. 5B) pointing to the fact the involved genes may play a role in cellular differentiation, cell proliferation and senescence regulation, phagocytosis and innate immunity. Among the top highly expressed genes, we found some small nuclear and long noncoding RNAs such as RN7SKP255, RN7SK, that are known to play a role in macrophage differentiation, polarization, and innate immune functions [32, 33]. IGHV1-69-2 gene encoding the Immunoglobulin heavy variable was also found among the top expressed genes alongside H2BP2, an extrachromosomal histone that stimulates innate and acquired immune defenses in different cells upon detection of double-stranded DNA (dsDNA) fragments derived from infectious agents or damaged cells [34, 35].

**Figure 5.**
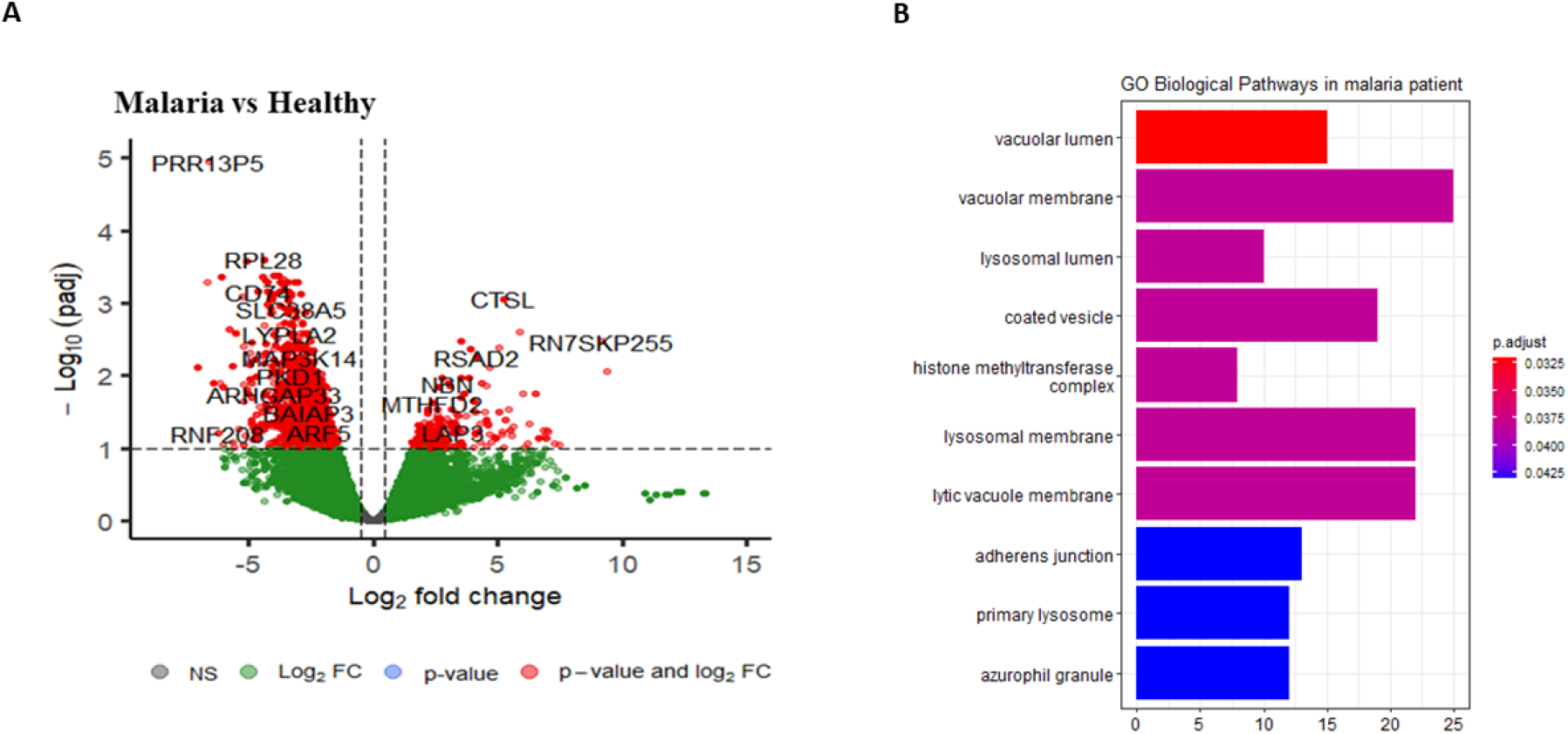
Volcano plot for comparison between malaria infected individuals and healthy. (A). The X-axis shows log fold change (positive values are up regulated relative to healthy. The Y-axis shows the −log10 of BH adjusted p-value (padj) value. The horizontal dashed line marks P= 1%, and the vertical dashed lines indicate two-fold expression difference among conditions. The differentially expressed genes are indicated in red (padj, 0.01 & log FC>1). Red points indicate upregulated genes, green points represent genes not passing the log2FC threshold but not the adj. p value, and grey points represent genes with no significant difference for both thresholds. B) GO-term functional enrichment in children with malaria infection. Bar plots showing nominal enrichment score.

### Transcriptional response between COVID-19 and malaria patients

Comparing gene expression in children with malaria to those with COVID19, we found about 2495 genes exclusively differentially expressed in the COVID-19 as compared to 1084 genes in the malaria group (Fig. 6B). About 62 genes were differentially expressed in both conditions. The majority (40/62) of these genes were upregulated in COVID-19 patients (Fig. 7) and could help us to determine which candidate genes to predicted for biomarkers from febrile patients either with malaria or COVID-19 (Table 5). Because COVID-19 and malaria both are infectious diseases which enhancing frequency of genes and pathways involved in the responses within the host. Some genes were also found to be expressed in the same direction. Three genes (ENSG00000234998, H2AC19 and TXNDC5) were both up-regulated in malaria and COVID. But the opposite was found with SIGLEC8, IL5RA and ALOX15 that were downregulated in both infections.

**Figure 6.**
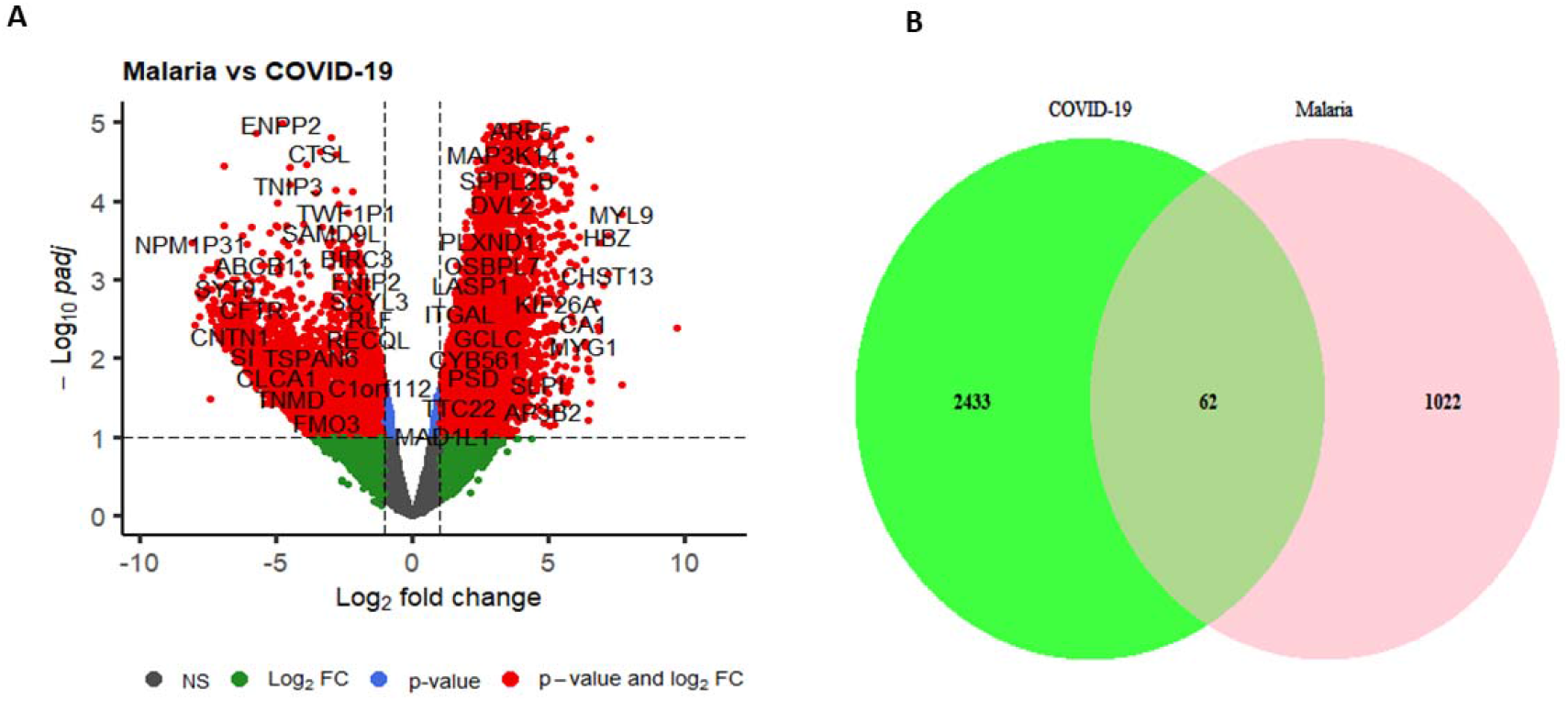
Immune system response in children with malaria infection and COVID-19 disease. **A**) Volcano plot for comparing between malaria and COVID-19 infected group samples. The X-axis shows log□ fold change (positive values are up regulated relative to malaria. The Y-axis shows the −log10 of BH adjusted p-value (padj) value. The horizontal dashed line marks P= 1%, and the vertical dashed lines indicate two-fold expression difference among conditions. The differentially expressed genes are indicated in red (padj < 0.01 & log□ FC>1). Red points indicate upregulated genes, green points represent without significantly different expression, and grey points represent genes with no significant difference. **B**) Venn diagram showing the shared, and unique numbers of differentially expressed genes in COVID-19 group and malaria group together. The overlaps of expression pattern of 62 genes differentially expressed were identified (padj< 0.1, |log2FC| > 1).

**Figure 7.**
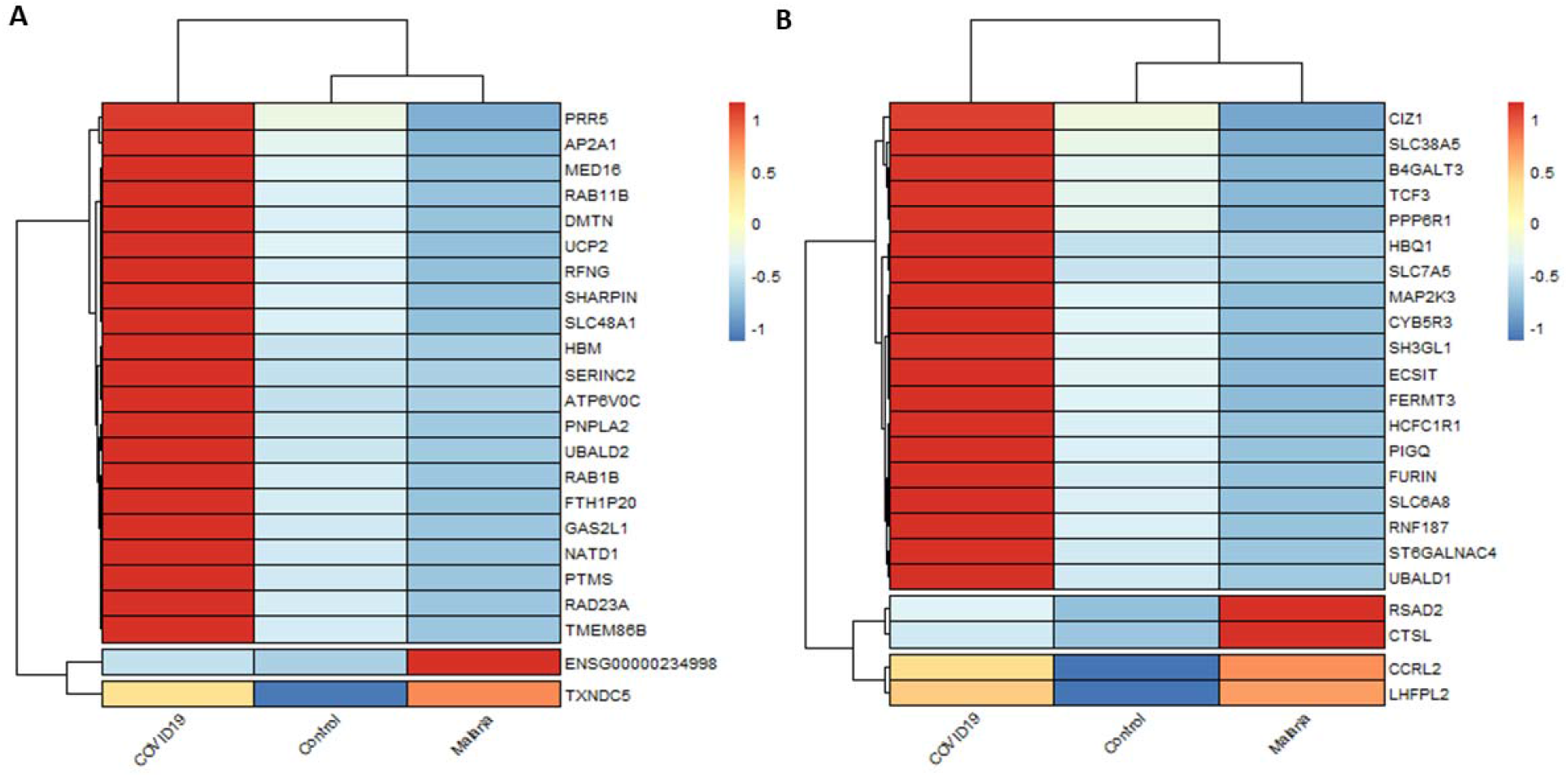
Representative target genes for diagnostics biomarker. Heat map of gene overlapped differentially expressed between malaria and COVID-19. The columns represent group of patients per infection and control as well. The rows indicate genes with significant differences in expression among both infections. The colors in the figure from red to blue indicate the level of gene expression from high to low based on z-score.

**Table 4.** List of top ten selected genes differentially expressed in children with malaria.

| Gene name | Description | logFC | padj |
| --- | --- | --- | --- |
| RN7SKP255 | RNA, 7SK Small Nuclear Pseudogene 255 | 9.1 | 0.0033 |
| IGHV1-69-2 | Immunoglobulin Heavy Variable | 7.4 | 0.0885 |
| FAM8A3P | Family With Sequence Similarity 8 | 7.0 | 0.0590 |
| RN7SKP203 | RNA, 7SK Small Nuclear Pseudogene 203 | 6.9 | 0.0567 |
| YBX1P6 | Y-Box Binding Protein 1 Pseudogene 6 | 6.9 | 0.0741 |
| SNORD71 | Small Nucleolar RNA, C/D Box 71 | 6.8 | 0.0726 |
| RNU6-485P | RNA, U6 Small Nuclear 485, Pseudogene | 6.8 | 0.0686 |
| H2BP2 | H2B Histone Pseudogene 2 | 6.6 | 0.0731 |
| ANKRD22 | Ankyrin Repeat Domain 22 | 6.4 | 0.0175 |
| CTSL | cathepsin L | 5.2 | 0.0008 |
| CD274 | CD274 Molecule | 4.5 | 0.0402 |
| GBP5 | Guanylate Binding Protein 5 | 4.0 | 0.0604 |
| GBP2 | Guanylate Binding Protein 2 | 2.9 | 0.0905 |

logFC: Log2fold change; padj: adjusted P value

**Table 5.** List of common genes differentially expressed in malaria and COVID-19

| Gene name | Malaria |  | COVID-19 |  |
| --- | --- | --- | --- | --- |
|  | log2 FC | padj | log2 FC | padj |
| **Upregulated (n = 3)** |  |  |  |  |
| ENSG00000234998 | 9.3 | 0.0085 | 6.8 | 0.0667 |
| H2AC19 | 5.8 | 0.0024 | 3.1 | 0.0667 |
| TXNDC5 | 4.8 | 0.0222 | 3.0 | 0.0717 |
| **Downregulated (n = 3)** |  |  |  |  |
| SIGLEC8 | -5.6 | 0.0074 | -3.1 | 0.0938 |
| IL5RA | -3.2 | 0.0397 | -3.1 | 0.0695 |
| ALOX15 | -5.4 | 0.0026 | -3.0 | 0.0766 |
| **Down and upregulated (n = 13)** |  |  |  |  |
| HBQ1 | -4.2 | 0.0163 | 4.1 | 0.0714 |
| HBM | -4.9 | 0.0070 | 3.7 | 0.0779 |
| SLC7A5 | -3.45 | 0.0051 | 5.7 | 0.0652 |
| SERINC2 | -2.9 | 0.0748 | 3.5 | 0.0719 |
| ATP6V0C | -3.1 | 0.0273 | 3.3 | 0.0712 |
| ST6GALNAC4 | -3.1 | 0.0099 | 3.0 | 0.0686 |
| RAD23A | -3.5 | 0.0055 | 2.9 | 0.0719 |
| PNPLA2 | -3.7 | 0.0010 | 2.8 | 0.0667 |
| GAS2L1 | -3.5 | 0.0173 | 2.8 | 0.0986 |
| TMEM86B | -2.8 | 0.0268 | 2.8 | 0.0715 |
| SLC6A8 | -3.0 | 0.0268 | 2.7 | 0.0999 |
| UBALD1 | -2.9 | 0.0187 | 2.6 | 0.0798 |
| RNF187 | -2.9 | 0.0121 | 2.5 | 0.0812 |

### Comparing GO term in malaria and COVID-19 patients

We performed GO (gene ontology) annotation analysis to explore the biological functions of the DEGs in malaria and COVID-19 group. Gene ontology analysis with the 62 common genes differentially expressed in malaria COVID-19 patients did not reveal any biological process or molecular function to be significantly enriched. However, cellular components such as hemoglobin complex and lipid particles were significantly enriched (Fig. 8). It is worth noting that those genes were to a large extent upregulated in COVID-19 as opposed to malaria patients.

**Figure 8.**
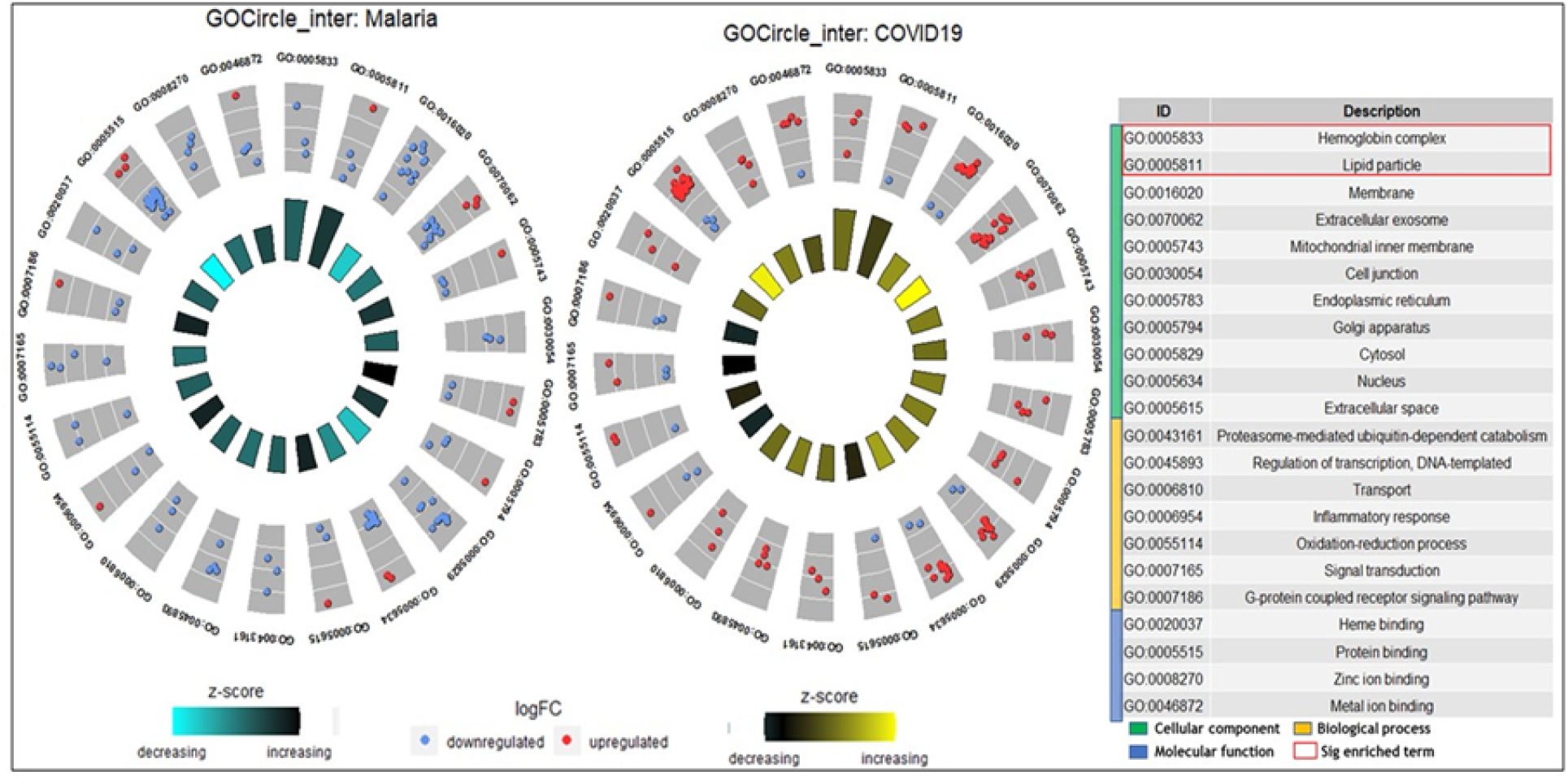
Gene ontology enrichment of overlapping gene (up-regulated and downregulated) between COVID-19 group and malaria group.

### K-means clustering associated with COVID-19 and malaria

To further confirm the differential gene expression between malaria and COVID-19 patients, K-means clustering was carried out. This analysis identified four main gene clusters (A-D) with cluster A and B genes having higher expression in COVID-19 patients as opposed to cluster C and D genes which had higher expression in malaria condition instead (Fig. 9A). Pathway analysis revealed that cluster A genes were associated with significant enrichment of immune-related biological processes such as neutrophil and granulocyte activation and degranulation, cell-mediated immune response and vesicle-mediated transport (Fig. 9B). Interestingly, cluster B genes were associated with hematopeosis, gas and oxygen transport confirming the gene ontology analysis done with the selected common 62 genes differentially expressed in malaria and COVID-19 conditions (Fig. 9B). Genes in cluster C are likely to favour innate immune processes, inflammation and immune regulation while Cluster D genes could be playing important role in mitotic cell division including chromatin assembly and chromosome organization (Fig. 9C).

**Figure 9.**
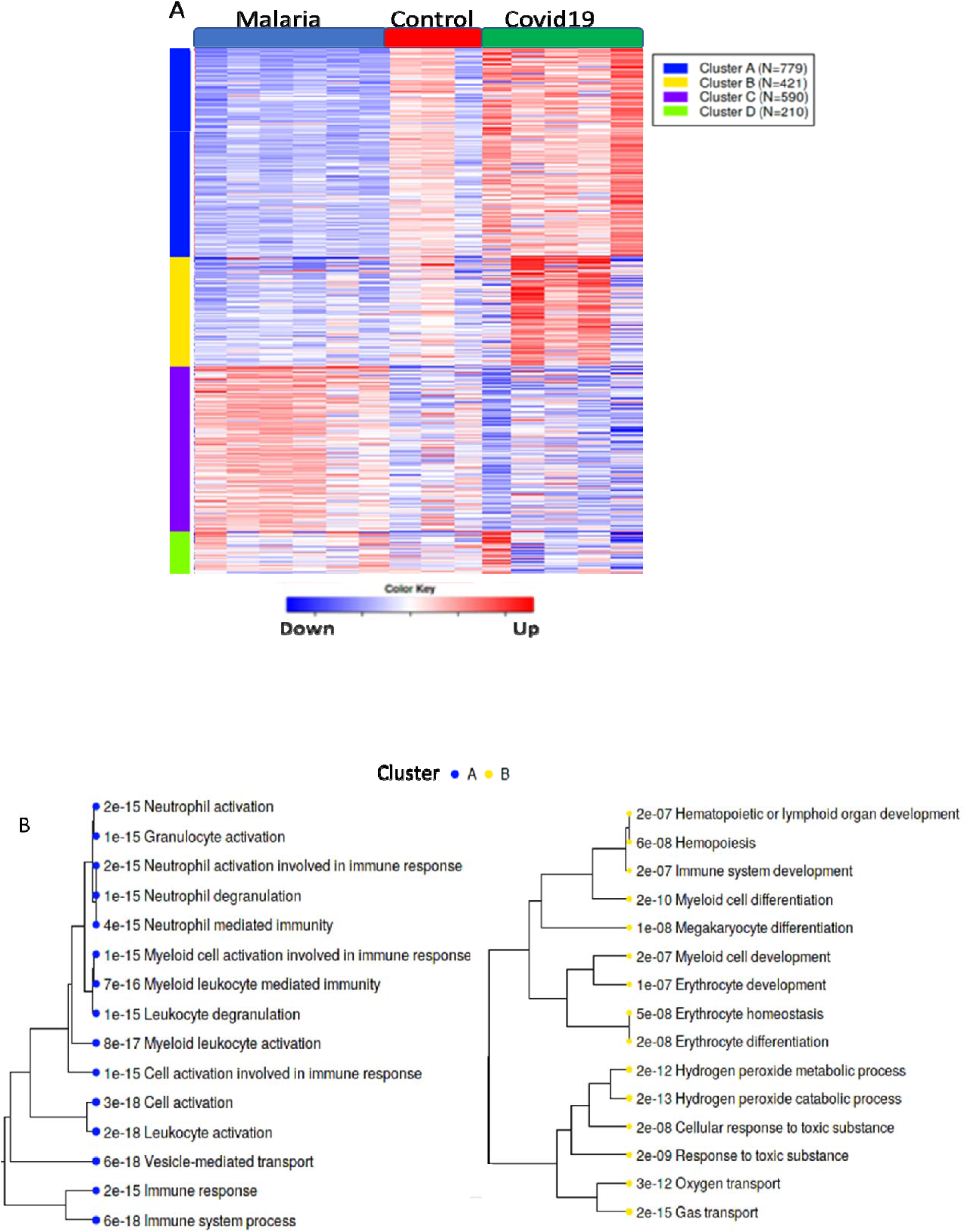
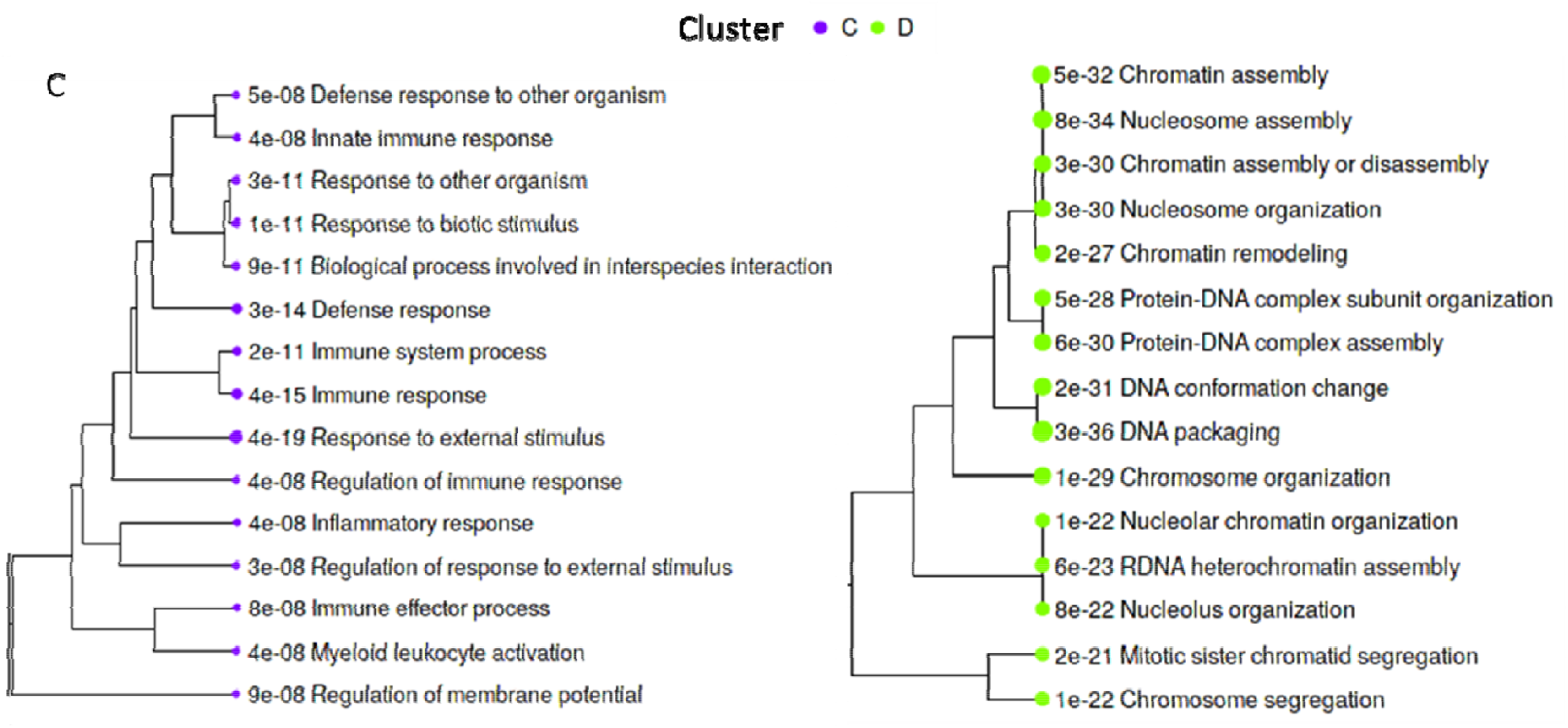
K-means clustering and associated pathway analysis. **A**) Heatmap showing clusters of genes differentially expressed in malaria, COVID-19 and healthy control. **B and C**) Pathway analysis, dendrograms showing biological processes significantly enriched per gene cluster. This analysis was performed on the Integrated Differential Expression and Pathway analysis (iDEP-95) platform.

## Discussion

In this study, we have used an RNA-seq approach to identify the transcriptional pattern of PBMCs/blood from infected children either with malaria or COVID-19 infection. This is useful because blood may be used to identify biological signatures within the hosts that interact with the pathogen and shorten the treatment [30]. As, we aimed to determine whether peripheral blood transcriptomic profiles are associated with the overlapping symptoms of malaria or COVID-19 infections in febrile children. This study has shown that differentially expressed genes are to a great extent linked to signaling and immune responses in COVID-19 and malaria conditions respectively. Commonly expressed genes were preferentially upregulated in COVID19 and downregulated in malaria making them potential reservoir of biomarkers to delineate the two conditions.

### Transcriptomic profile of children with COVID-19 and healthy group

In children with COVID-19, a total of 2495 gene were differential expressed (188 upregulated and 2307 downregulated) and the main biological pathways were associated to the regulation of membrane potential and signaling (Fig. 4B). Among genes involved in positive regulation of cellular process we found (TCAF2C, AchE, CD151), whereas TCAF2C is TRPM8 channel associated factor 2C and an intracellular gene with unknown function but it predicted to initiate transporter binding activity. We observed also the presence of AchE gene which can be useful to monitoring the cholinergic system in COVID-19 patients [36] and CD151 gene which participate during endocytosis mechanism of the viral entry [37]. Among the genes participate in positive regulation of biological process we also found (C1QC, TCAF2C, NECTIN2) and among genes initiate the cellular catabolic process we identified (ACHE, and UBE2C) (Table 3). Notably, the comparison of COVID-19 samples relative to healthy samples, we found that about 188 upregulated genes differentially expressed with SLC12A5-AS1, ENSG00000234998, GPS2P1, RCCD1-AS1 and RAP1GAP being among the top expressed genes (Fig. 4). Some of the top DEGs have previously been shown to associated with viral infections. These include SLC12A5-AS1 which is a lncRNA gene associated with NK-cell in the lung and can respond to viral infections [38]. We also found the upregulation of ENSG00000234998 as a novel transcript, an antisense lncRNA gene to FCGR1B, and is associated with CD64, which is the receptors of SARS-CoV-2 [39]. These receptors have been reported as biomarkers for early viral infections. We detected a significant expression of RAP1GAP (GTPase activating protein) which is involved in the viral invasion through the change in the epithelial morphogenesis and intercellular tight junction formation [40] (Table 3). The CD151 molecule another upregulated gene has been shown to be associated with asthma thereby inducing airways hyperresponsiveness via calcium signaling. Difficulty in breathing is one of the complications of COVID-19 and this may explain the upregulation of CD151. Moreover, this has also been shown to be involved in influenza leading to increased disease severity [30].

### Comparison of children with malaria and healthy group

We compared the malaria samples relative to the healthy samples, and identified about 1084 genes differentially expressed (179 upregulated and 905 down regulated) with ENSG00000234998, RN7SKP255, IGHV1-69-2 and FAM8A3P falling among the top five highly expressed in children with malaria (Fig. 5). In the children with malaria, we identified the enhancement of an anti-bacterial gene like IFI44 gene and many other genes associated with immune response processes (Table 4). Genes such as RN7SKP255, RN7SKP203 and RN7SK contain 7SK as a molecule participate in controlling multiprotein complex P-TEFb (the positive transcription elongation factor) [41]. IGHV1-69 is an immunoglobulin heavy variable 1-69-2, reported to be used by germline gene to neutralizing antibodies [42]. RPL34 (60S ribosomal protein L34) belong to ribosomal proteins in large subunits, and it plays a key role in cell cycle distribution, cell proliferation and apoptosis [43]. FAM8A3P (a family with sequence similarity 8, member A3 pseudogene), the expression level of ANKRD22 (Ankyrin Repeat Domain-Containing Protein 22) involved in cell proliferation and enhanced the glycolysis associated with an increase in AMP/ATP and decrease in ATP/ADP levels [44]. ANKRD22 downregulation has been shown to inhibit an excessive inflammatory response [45]. The CD274 gene is involved in adaptive immune response by binding to PD-1 for inhibitory signaling, which inactivates immune cells such as is expressed on activated B cells, T cells, and NKT cells, which are crucial in homeostasis of immune systems during infections [46]. The GBP5 gene involved in AIM2 inflammasome activation in response to bacterium where it protect host against pathogen [47]. In parasites infection, the GBP2 gene is involved in nuclear mRNA export and it promotes the asexual and sexual development of Plasmodium [48]. H2BP2 (Histone Cluster 2 H2B) mostly involved in nucleosome structure. It acts as a cytoplasmic sensor to identify dsDNA debris derived from infectious agents or damaged cells with subsequent activation of activate innate and acquired immune responses in various cell types resulting in dsDNA-induced type I IFN production [34]. We also identified various genes of unknown function among the top up-regulated genes in children with malaria. We identified hemoglobin subunits to be upregulated in children with COVID-19 (Fig. 9). Hemoglobin is a key play in oxygen transport in the system. CYB5R3 a cytochrome b5 reductase 3, which transfers electrons from NADH to cytochrome b5 in red blood cells, was also found upregulated in children with COVID-19. In malaria, the mutation of CYB5R3 has been shown to be associated to the risk of developing severe anemia in children [49, 50]. The high expression of RSAD2 gene, an interferon-inducible gene, reported to be involved in the innate immune response against viruses [51] was upregulated in both malaria and COVID-19 conditions. This could be due to the intracellular nature of both pathogens. The enzyme Cathepsin L (CTSL) an endosomal cysteine protease cleaves the spike protein of SARS-CoV-2 and enhanced virus entry into the host [52].

### Overlapped genes are more upregulated in children with COVID-19 than in those with malaria

Three genes, ENSG00000234998 (a lncRNA class), H2AC19 and TXNDC5 were found upregulated in malaria and COVID-19 infections. Conversely, SIGLEC8, IL5RA, and ALOX15 were expressed at lower levels in in both infections compared to the healthy as control. H2AC19 gene belongs to the histone H2A family. It plays a crucial role in DNA and protein binding [53]. TXNDC5 gene inhibits apoptosis. It is involved in pathogenesis, proper protein folding and cell proliferation [54]. SIGLEC8 is a protein coding gene belonging to the CD33-like subgroup of SIGLECs, involved in plasma membrane and innate immune response. PGD2 receptors also known as CD294 is expressed in CD4+ effector Th2 cells (T helper 2) cells [55]. Previous studies suggested that PGD2 receptor is mediated in aspirin-induced respiratory disease like COVID-19 [56]. ALOX15 gene belongs to lipoxygenases enzymes and expressed in immune cells. In addition, ALOX15 is more localized in plasma membrane and at the cytoplasmic side of intracellular membranes [57]. The IL5RA gene binds to interleukin-5 (IL5) and mediate the activation of the transcription factor SOX4 [58]. Chrisopher *et al*. in their findings showed that the presence of eosinophil activity in COVID-19 patients. When the eosinophil activity is high indicates a parasitic infection because participates to both adaptive immunity and innate [59, 60]. Therefore, IL5RA due to the activity of eosinophils associated with IL5 makes it a potential therapeutic target.

### Transcriptional signatures associate with febrile patient of malaria and COVID-19

Overall, we identified 3583 differentially expressed genes (1084 malaria and 2495 in COVID-19) and among of them about 62 genes were common differentially expressed in both infections (Fig. 6). The common genes in immunologically relevant molecular signatures between a patient with either malaria or COVID-19. About 13 genes were found to be downregulated in children with malaria relative to healthy baseline, and then were uniquely expressed at higher levels in children with COVID-19 infection (Table 3). Among these genes, HBQ1 has been reported to participate in O_2_ transport in human fetal erythroid tissue, expressed in neutrophils [61]. HBM gene is expressed in neutrophils and involved in humoral immune response and O2 transport and [62]. The SLC7A5 gene (Solute Carrier Family 7 Member 5) is an amino acids transmembrane transporter, like leucine, glutamine and tryptophan, which are key amino acids involved in cellular function and metabolic activation [63]. We found that the expression of SLC7A5 (LAT1) is higher in COVID-19 patients and lower in malaria patients. It has been postulated that SLC7A5 gene is associated with transport of essential amino acids in pathogenesis [64, 65]. Its expression has been reported to be mediated by cytokine IL-2 in malaria patients [64, 65] and occurs in the earlier host response against Plasmodium infection eg; *Plasmodium yoelii* [66]. The SERINC2 gene (Serine incorporator 2) belongs to SERINC family and participates in the transmembrane proteins formation by incorporating serine residues into membrane lipids. Strikingly, we found that SERINC2 gene was significantly upregulated in COVID-19 and downregulated in malaria patients. This observation is in line with a previous report which showed high expression of SERINC2 gene in COVID-19 patients [67] (Table 5). The ATP6V0C (ATPase H+ Transporting V0 Subunit C) is a component of V-ATPase (vacuolar ATPase) which inhibits lysosomal acidification mechanism by blocking the passage of H^+^ via lysosomal membrane [68] and it has been found that S protein of SARS-CoV-2 induces the higher expression of V-ATPase within COVID-19 patients [69].

The RAD23A gene is a nucleotide excision repair protein, we found that it is downregulated in malaria patient. Indeed, this was similar to the a published report that showed low RAD23A expression in children with malaria infection [70]. PNPLA2 gene, contributes in lipid metabolism [71]. The GAS2L1 (growth-arrest-specific 2 (GAS2)-like 1) gene, is a member of the GAS2 family widely expressed in human tissues. In addition, it plays a major role in centrosome motility and the overexpression of GAS2L1 induces the centrosome disjunction but the mechanism still unclear [72]. TMEM86B (transmembrane protein 86B) is a member of the YhhN family, which is involved in lipid metabolism with probable modulation of the level of plasmalogen in cell lines [73]. SLC6A8 (Solute Carrier Family 6 Member 8), a creatine transporter located in cell membrane and macrophages use SLC6A8 gene to accumulate high level of intracellular creatine [74]. UBALD1 gene (UBA Like Domain Containing 1) also was found to be upregulated in peripheral blood mononuclear cells collected from COVID-19 patients [75, 76]. We noted, a similar upregulation in children with COVID-19 infection. RNF187 gene (E3 ubiquitin-protein ligase RNF187) is involved in transfer of ubiquitin and innate immune response. This gene has been reported to be expressed in platelets from COVID-19 patients [77, 78] as observed in the present study. We also noticed that among the shared genes, the hemoglobin complexes and lipid mediators are differentially expressed between the COVID-19 samples and malaria samples (Fig. 8). Thus, we suggest that HBQ1, and HBM together with the common genes described here may constitute good candidates for the development of a diagnosis biomarker to delineate of malaria from COVID-19 infections as they have contrasting expression levels in both infections.

### Gene ontology revealed that inflammation and innate immunity to be enriched in children with both COVID-19 and malaria infections

The K-means clustering algorithm and GO analysis identified three main biological pathways to be enriched such as cell-mediated immunity, inflammation and innate immunity in children with both COVID-19 and malaria infections. The Corona virus and Plasmodium parasite are both intracellular pathogens which are mainly fought by the cell-mediated arm of the immune system, including the innate branch [79–81]. Biological processes associated with hematopoiesis, oxygen and gas transport were found significantly enriched in COVID-19 patients. COVID-19 disease is known to be associated with cough and difficult breathing which may have as corollary, low oxygen levels in blood and tissue hence, the enhancement of hematopoietic processes and gas and oxygen transport. This could be supported further by the upregulation of CD151 known to induce increased disease severity in asthma and influenza infections[30]. Among the common genes highly differentially expressed (Fig. 9). Using gene set enrichment analysis, we tested whether the upregulated genes either from COVID-19 or malaria group are discriminatory in GO terms and we found that children with COVID-19 induce more genes involved regulation of biological and catabolic process.

## Conclusion

We scrutinized the transcriptional responses in children with malaria and children with COVID-19 infection. Genes with contrasting expression levels in both conditions were identified and could be used in the development of diagnostic or prognostic markers for those infections. These could help speed up decision-making on the treatment of febrile patients with suspected malaria with COVID-19 co-infection. We recommended further research with coinfection samples which could help identify potential co-infection markers particularly in regions with high malaria prevalence.

### Limitations of the study and suggestions for further work

The predicted role of different genes (upregulated and downregulated) needs to be further investigated either in children or adults with data generated from scratch instead of using RNA seq data from public repositories which are produced in different high-throughput sequencing methods for diverse scientific goals. Another limitation was removing batch effect in RNA seq data and the missing data of coinfection case in children. This would have enabled us to generate distinctive transcriptomic profiles of responses in patient with malaria and COVID-19 disease as coinfection.

## Data acquisition and software availability

The RNA-seq data were collected from public databases to study children infected with malaria and COVID-19 infection. Only datasets with paired end reads were used in this study. The study examined the transcription profile responses of children with *Plasmodium falciparum* and severe acute respiratory syndrome coronavirus 2 (SARS-CoV-2). The datasets used for children with malaria are available on European Nucleotide Archive (ENA) through the study accession number PRJEB33892. The datasets used for children with COVID-19 infection are available on Gene Expression Omnibus (GEO) through accession number GSE178388. The reference genome used for indexing and mapping was downloaded from <u>GENCODE</u>. The annotation reference genome used for counting reads was downloaded from <u>ENSEMBL</u> genome database. The data files for this study, as well as the source code, Software tools, databases and command line used for preprocessing data under Linux and R scripts for downstream analysis, are available on Github. (https://github.com/omicscodeathon/RNA-seq-Malaria).

## Author contributions

Conceptualization, <u>N.L</u>; J.A.K.O; Formal analysis, <u>N.L</u>; J.A.K.O; Investigation, <u>N.</u>L; J.A. K.O; W.M. R; K.M; U.<u>D;</u> P.K. Methodology, <u>N.L</u>; J.A.K.O. Resources, O.I.A; N.L. Data Curation, <u>N.L</u>; J.A.K.O. Project Administration, O.I.A. Resources, O.I.A; N.L. Supervision, O.I.A; <u>N.L</u>. Writing and original Draft Preparation, W.M.R; U.D; N.L; Writing–Review & Editing, N.L; J.A.K.O; W.M.R; K.M; U.D; P.K; O.I.A. Funding Acquisition, O.I.A; A.D. All authors have read and agreed to the published version of the manuscript.

## Acknowledgements

This study was funded by the National Institutes of Health Office of Data Science Strategy (ODSS), in partnership with the African Society for Bioinformatics and Computational Biology (ASBCB). We are grateful for their support to provide High performance computing (HPC) in the cloud for RNA seq data analysis. Centre for Research in Infectious Diseases (CRID), for the onsite technical support.

